# Convergent mechanisms, divergent strategies: a comparison of nectar intake between a generalist and a specialized bat species

**DOI:** 10.1101/2024.11.28.625920

**Authors:** Laura L. Quinche, Felipe Garzón-Agudelo, Sharlene E. Santana, Hugo F. López-Arévalo, Alejandro Rico-Guevara

**Author notes:** Corresponding author: Laura L. Quinche;.

## Abstract

Nectar-feeding bats exhibit a range of specialized adaptations that allow them to efficiently extract nectar from flowers. These adaptations include diverse tongue morphological traits and feeding strategies that reflect varying degrees of specialization for nectarivory. While the feeding mechanisms of highly specialized nectar-feeding bats are well-studied, little is known about the feeding behaviors of non-specialized species like *Phyllostomus discolor*. This study compares the nectar extraction behaviors of *P. discolor* and the specialized *Anoura geoffroyi*, examining morphological and biomechanical adaptations that affect nectar-feeding efficiency and foraging strategies. Using high-speed videography, we analyzed the feeding behaviors of both species, focusing on tongue kinematics, and feeding efficiency. Both species used a brush-tongue lapping technique but exhibited notable behavioral and kinematic differences, resulting in efficiency variations. *P. discolor* has a shorter, less flexible tongue than *A. geoffroyi*, though its tongue shows similar mobility capacities (licking frequency). Unlike *A. geoffroyi*, which hovers to feed, *P. discolor* lands, allowing for longer visits and greater nectar extraction per visit. However, *P. discolor* demonstrated lower feeding efficiency, likely due to its reduced tongue specialization for nectarivory. These findings reveal convergence in the general feeding mechanism but highlight differences in morphological and behavioral specialization that affect feeding kinematics and efficiency. Our study illuminates how foraging strategy and tongue morphology impact feeding efficiency, pointing to evolutionary pathways that promote niche differentiation within nectar- feeding bat communities.

## INTRODUCTION

Nectar is one of the most ubiquitous food resources on Earth and it mediates mutualistic relationships between plants and animals (Nicolson et al., 2007). Flower visitors consume this highly nutritious reward produced by plants and participate in plant pollination by transporting and transferring pollen (Nicolson et al., 2007; Parachnowitsch et al., 2019). There is a wide diversity of nectar consumers, from jumping spiders to bats (Fleming and Muchhala, 2008; Jackson et al., 2001), with varying degrees of nectarivory that span an ecological continuum from opportunistic to highly specialized flower visitors.

Within the diverse order of bats (Chiroptera), the nectarivorous diet has evolved independently in Paleotropical fruit bats (Pteropodidae), Neotropical leaf-nosed bats (Phyllostomidae) (Fleming et al., 2009), and in two species of the families Vespertilionidae (*Antrozous pallidus*, Frick et al., 2014) and Mystacinidae (*Mystacina tuberculata*, Arkins et al., 1999), respectively. Within Phyllostomidae, nectarivory evolved independently as the main feeding habit in three lineages: the Glossophaginae and Lonchophyllinae subfamilies and the *Phyllostomus* genus (Rojas et al., 2011). Although nectarivory has been recognized as the main feeding habit in these three lineages, they have different degrees of dependence on nectar. Most species in Glossophaginae and Lonchophyllinae are specialized nectarivores, meaning they exhibit behavioral, morphological, and physiological adaptations to this diet (Gonzalez-Terrazas et al., 2012). Species of the genus *Phyllostomus* are omnivorous (Gardner, 2008), and *Phyllostomus discolor* Wagner 1843, exhibits a high level of nectarivory in their diet (82.1% of pollen and nectar; Kalko et al., 1996) or evidence of floral visitation (70.4% of pollen in total diet; Giannini and Kalko, 2004).

A bat extracts nectar from a flower by bringing its snout close to the corolla and extending its tongue into the nectar chamber; thereby the tongue becomes the primary food acquisition organ. Researchers have sought to elucidate the importance of the tongue during feeding in specialized nectarivorous bats by characterizing tongue morphology (Harper et al., 2013; Nicolay and Winter, 2006) and nectar-feeding behavior (Bechler et al., 2024; Gonzalez-Terrazas et al., 2012; Nicolay and Winter, 2006; Tschapka et al., 2015) and relating these traits to foraging efficiency. In specialized nectarivorous bats, two different morphological and biomechanical strategies have been observed for extracting nectar from flowers. *Glossophaga soricina*, a glossophagine species, have hair-like (filiform) papillae on its tongue and employs a lapping mechanism in which the papillae are erected by blood flow (brush-tongue lapping technique) (Harper et al., 2013; Tschapka et al., 2015). In contrast, *Lonchophylla robusta*, a lonchophylline species, has lateral grooves in the tongue, and its extraction mechanism consists of pumping nectar into the mouth through the lingual grooves (pumping-tongue drinking technique) (Tschapka et al., 2015). Little is known about the morphology associated with nectarivory in non-specialized species (e.g., *Phyllostomus* spp.; Nicolay, 2001; Nicolay and Winter, 2006; Quinche et al., 2023) and the nectar extraction mechanism of non-specialized nectarivorous species has not been described. Examining the morphology and nectar uptake behavior of non-specialized species is essential to determine whether additional modes of nectar extraction have evolved in bats and could shed light on the initial steps to specialized nectarivory in the lineage.

Morphological and behavioral traits related to nectar consumption have been recognized in *P. discolor*, such as a long and highly mobile tongue with hair-like papillae on its surface (Quinche et al., 2023), and a nectar extraction rate as efficient as that of *Glossophaga soricina* (Nicolay, 2001; Nicolay and Winter, 2006). However, a complete description and a quantitative study of *P. discolor* nectar uptake behavior and a comparison with specialized species is lacking. Filling this knowledge gap is important to better understand the relationship between feeding behavior and efficiency in species with different degrees of morphological and ecological specialization. In this study, we conducted comparative experiments between the non-specialized *P. discolor* and *Anoura geoffroyi* Gray 1838, a nectar specialist glossophagine. Previous studies have described morphological adaptations to nectarivory in *A. geoffroyi*, such as a highly protrusible tongue with hair-like papillae (Bhatnagar and Smith, 2007; Muchhala et al., 2024). However, no research has been conducted to describe the extraction mechanism and nectar-feeding efficiency of this specialized species.

The extraction mechanism employed by the genus *Phyllostomus* is not yet known. We hypothesized that the nectar extraction mechanism of *P. discolor* resembles the brush-tongue lapping technique of Glossophaginae species for two main reasons. First, from an evolutionary perspective, we would expect lapping to be the first stage in nectarivory, as it resembles the use of the tongue to drink water in mammals with incomplete cheeks (Williams, 2019); more elaborate mechanisms, such as those seen in Lonchophyllinae, would have evolved later in response to particular selective pressures during plant-pollinator interactions (e.g., evolution of long corollas and tongues that can extract the nectar deep inside without spending extra time reciprocating). Second, from a morphological perspective, we would expect *P. discolor* to employ its tongue similar to glossophagines, as both taxa share hair-like lingual papillae, differing from Lonchophyllines as *P. discolor* lacks groove-like structures on the sides of the tongue.

Given the considerable differences in size (*P. discolor* 39.7 - 44.6 gr. and *A. geoffroyi* 8.5 - 13.0 gr., Kwiecinski, 2006; Oprea et al., 2009), foraging strategy, diet specialization, and evolutionary history, we predicted that even though the mechanism could be similar (brush-tongue lapping technique), there would be clear dissimilarities in nectar feeding behavior between *P. discolor* and *A. geoffroyi*. Specifically, we expected *P. discolor* to exhibit longer visits as it lands to feed (Giannini and Brenes, 2001; Pedrozo et al., 2018), as opposed to *A. geoffroyi*, which would hover while feeding and therefore would also be limited by the energy invested in hovering ((Voigt, 2004; Voigt and Winter, 1999). Additionally, we predicted that the licking frequency, amount of nectar extracted, and drinking efficiency would be higher in *A. geoffroy*i because of its more specialized feeding habits and tongue morphology. Using high-speed videos, we described and compared the nectar-feeding behavior of *P. discolor* and *A. geoffroyi* by characterizing their foraging behaviors and nectar extraction mechanisms and by measuring the nectar extracted, time of visit, extraction efficiency, maximum tongue protrusion, and licking frequency of both species.

## MATERIALS AND METHODS

### Field survey, captivity, and specimen collection

Fieldwork was conducted at two different locations and periods: (1) in Hacienda La Cabaña (Sede Cabaña), Cumaral, Meta, Colombia between May and June 2019, and (2) in Colibrí Gorriazul Research Center, Fusagasugá, Cundinamarca, Colombia, in October 2019, February 2021, and January 2023. Bats were captured with mist nets, and all bats were released, except for adult *Phyllostomus discolor* (n=10) and *Anoura geoffroyi* (n=11) individuals, which were held in captivity in a flight cage (2.5 × 2.5 × 2.5 m). Female bats were confirmed to be neither pregnant nor nursing. Between both species, we kept a maximum of four individuals in the cage at the same time. We did not see any aggressive interactions among individuals while sharing the cage during the experiments. In the cage, the animals could fly freely and hang from the walls and ceiling. We placed one or two feeders designed like a banana inflorescence (a common local food source known to be visited by both species), with a hemispherical cone shape approximately 30 cm long, which allowed the bats to hang while drinking nectar as needed. We attached a transparent flat-sided container (test container: 11 mm × 11 mm × 44 mm) to one of the feeders.

After capture, the bats were habituated to the experimental setup inside the cage (test container and cameras/lights) by first recording videos of them drinking nectar while holding them in our hands (hand-held videos). The position of the feeder was kept constant, and it was shown to the bats by releasing them above the feeder. The bats were trained for one night, after which all of them learned to visit the test container by flying to it on their own and returning to their preferred hanging spot.

During captivity, the bats were fed artificial nectar, fruit, and water *ad libitum* while in the cage. Every one or two nights bats were taken from the cage to individually feed them a mixture of artificial nectar with crushed mealworms before being released back into the cage. Captured individuals spent 2–4 nights in captivity, after which they were released at the capture site. We collected two adult, non-pregnant, non-lactating female individuals as voucher specimens: one of *P. discolor* during fieldwork in Cumaral, Meta, Colombia, and one of *A. geoffroyi* during fieldwork in Fusagasugá, Cundinamarca, Colombia. The individuals were held in captivity and collected under the institutional collection permits of the Universidad Nacional de Colombia (number: 0255, March 2014). We followed the IACUC guidelines recommended by the American Society of Mammalogists for all procedures (Sikes and the Animal Care and Use Committee of the American Society of Mammalogists, 2016).

### Tongue morphology

To describe and compare the lingual surface of the two species, we dissected the tongues from the two specimens collected, and prepared them for scanning electron microscopy (SEM). First, we fixed the tongues in 4% formaldehyde, dehydrated the samples, and covered them with a gold coating. We obtained the SEM images from the Laboratorio de Microscopía Electrónica de Barrido at the Universidad Nacional de Colombia with an FEI Quanta 200 (FEI Company, Eindhoven, the Netherlands) and from the Laboratorio de Microscopía Electrónica de Barrido at the Pontificia Universidad Javeriana de Colombia with an EVO HD15 (Zeiss, Jena, Germany), these scanning electron microscopes were operated at an accelerating voltage of 30 and 10 kV and direct magnifications up to 1,000x.

### Nectar extraction experiment design

Bats were recorded visiting the test container filled with artificial nectar (sucrose solution at 17% w/w concentration) at different distances from the upper rim (i.e., nectar depth). *P. discolor* and *A. geoffroyi* individuals were filmed with high-speed video cameras (JVC GC-PX100BU, Fastec TS5, and Chronos 1.4, Krontech) set at 240 – 1000 frames per second. We made videos with two main lighting configurations: LED light at the back or side of the tube and infrared (IR) lights from the outside of the feeder setup and aimed at the feeder. We focused the camera on the test container using tele-macro modes/lenses. We recorded two types of high-speed videos: 1) videos of hand-held individuals drinking nectar in different views (lateral, frontal, and ventral; hand-held videos) and 2) videos of free-flying individuals inside the cage approaching volitionally to drink nectar (free-flying videos).

### Video data collection

We used ImageJ-Fiji (v2.14.0; National Institutes of Health, Bethesda, MD, USA) to quantify variables from high-speed videos. To characterize tongue movements, we tracked the tongue tip movements of nine visits of *P. discolor* and 17 visits of *A. geoffroyi* while feeding from the test container at nectar depths ranging from 10 to 15 mm. Additionally, to facilitate comparisons between species, we analyzed the tracking data of the first lick from each visit. Since the number of licks and their durations varied in every visit (within species), we normalized the lick duration to a percentage scale, with 0% marking the start and 100% the end of the lick. We then calculated and plotted the mean first lick pattern for each species.

To characterize the general drinking behavior of both species (beyond tongue kinematics), we used videos of 40 visits of *P. discolor* and 88 visits of *A. geoffroyi* feeding at a wider range of nectar depths (from 2 to 30 mm). We measured complete visits, defined from the first appearance of the tongue tip (start of the visit) until the moment the tongue was entirely retracted into the mouth (end of the visit), regardless of the total number of licks. In all of these visits, the tongue contacted the liquid at each lick. We measured the snout-tip and tongue-tip insertion, recording how far the bats inserted their snout and tongue tip into the test container (in mm). The amount of nectar extracted (g) was calculated by measuring the change in the meniscus of the nectar in the test container before and after the visit, then calculating the volume using the dimensions of the test container, and by using the density of the sucrose solution provided. Visit time (s) was measured from the first frame where the tongue appeared to the last frame where it was no longer visible as the bat left the nectar test container. This frame count was then converted to seconds according to the frame rate. We calculated nectar extraction efficiency, *E* (g s^-1^), by dividing the amount of nectar extracted by the total visit time. Then, for a more meaningfulcomparison between species of different sizes, which will reflect their different energy demands, we calculated the size-adjusted efficiency *Es* (g s^-1^ g^-1^) as follows:

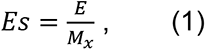

where *M_x_* is the body mass of species *x* in grams.

The mean maximum tongue protrusion length (mm) was measured from the snout tip to the tongue tip along the dorsal surface of the tongue. This measurement was taken from lateral videos during maximum tongue protrusion events, defined as at least three consecutive attempts to reach deeper nectar without success and with minimal change in tongue extension. Notably, maximum tongue protrusion differs from tongue-tip insertion and maximum extraction depth. Maximum tongue protrusion follows the curved surface of the tongue, while tongue-tip insertion and maximum extraction depth are measured as straight distances. Tongue-tip insertion refers to the distance the tongue tip extends inside the tube, whereas maximum extraction depth indicates the deepest nectar level reached by the tongue. Finally, we calculated licking frequency (licks s-1) by dividing the number of licks by the visit time.

### Statistical analysis

We used Generalized Linear Models (GLMs) to explore the effects of species and nectar depth as explanatory variables on various response variables that describe drinking behavior. The response variables were: the number of licks, amount of nectar extracted, visit time, nectar extraction efficiency, standardized extraction efficiency, lick frequency and maximum tongue extension. We employed the Poisson distribution and log link function for the dependent variable number of licks and the Gamma distribution and the inverse link function for the rest of the dependent variables. Statistical tests were performed using R (v4.2.3; R Core Team 2023).

## RESULTS

### Overall behavior during feeding

Once the bats were released into the cage, after being hand- held fed at the feeder with the test container, they flew in circles and explored the space before approaching the test container. All captive Individuals learned the location of the nectar source and began visiting the feeders on their own on the first or second night. *Phyllostomus discolor* always landed on the feeders before drinking. First, they landed on different parts of the feeder to then crawl to approach the nectar container, and after some visits, they landed directly in front of the container. Once they found the feeder, they landed on top of it and hung upside down, sometimes keeping their wings open and other times folding them while drinking nectar. Conversely, *Anoura geoffroyi* always approached the test container and drank nectar while hovering, making quick, consecutive visits.

Although *P. discolor* individuals usually drank nectar until they seemingly could not drink any more (because they could not reach deeper in the container), *A. geoffroyi* individuals spent almost the same time during each visit, reaching deeper but without emptying the test container. Both species rarely closed their eyes while feeding, and no territorial behavior was observed around the feeders. When more than one individual of the same species was in the cage, sometimes the bats left their roosting sites and foraged together, taking turns to access the nectar. We observed this joint feeding behavior more in *A. geoffroyi* than in *P. discolor*.

### Tongue kinematics and morphology

The mechanism employed by *P. discolor* and *A. geoffroyi* consisted of repetitive tongue protrusion and retraction cycles, a lapping mechanism (e.g., Figs. 1A,B, 2A, Movies 1, 2 in supplementary material). Once *P. discolor* started feeding, it protruded its tongue and dipped it into the nectar stretched and flattened dorsoventrally, in a curved ventral motion. At maximum extension on each lick, the flattened tongue folds inwards with the raised edges forming a partial channel medially, and the tongue tip further curves caudally. This curvature creates a kind of hook on the ventral side of the tongue tip (Fig. S1). However, when the nectar level was low and the individual attempted to reach it, the tongue did not bend downwards and backwards as in higher nectar levels. Similar tongue bending was observed in *A. geoffroyi* individuals; however, their tongues were more mobile and bent in multiple directions and with less consistency (e.g., Fig. 1B,D).

**Fig. 1.**
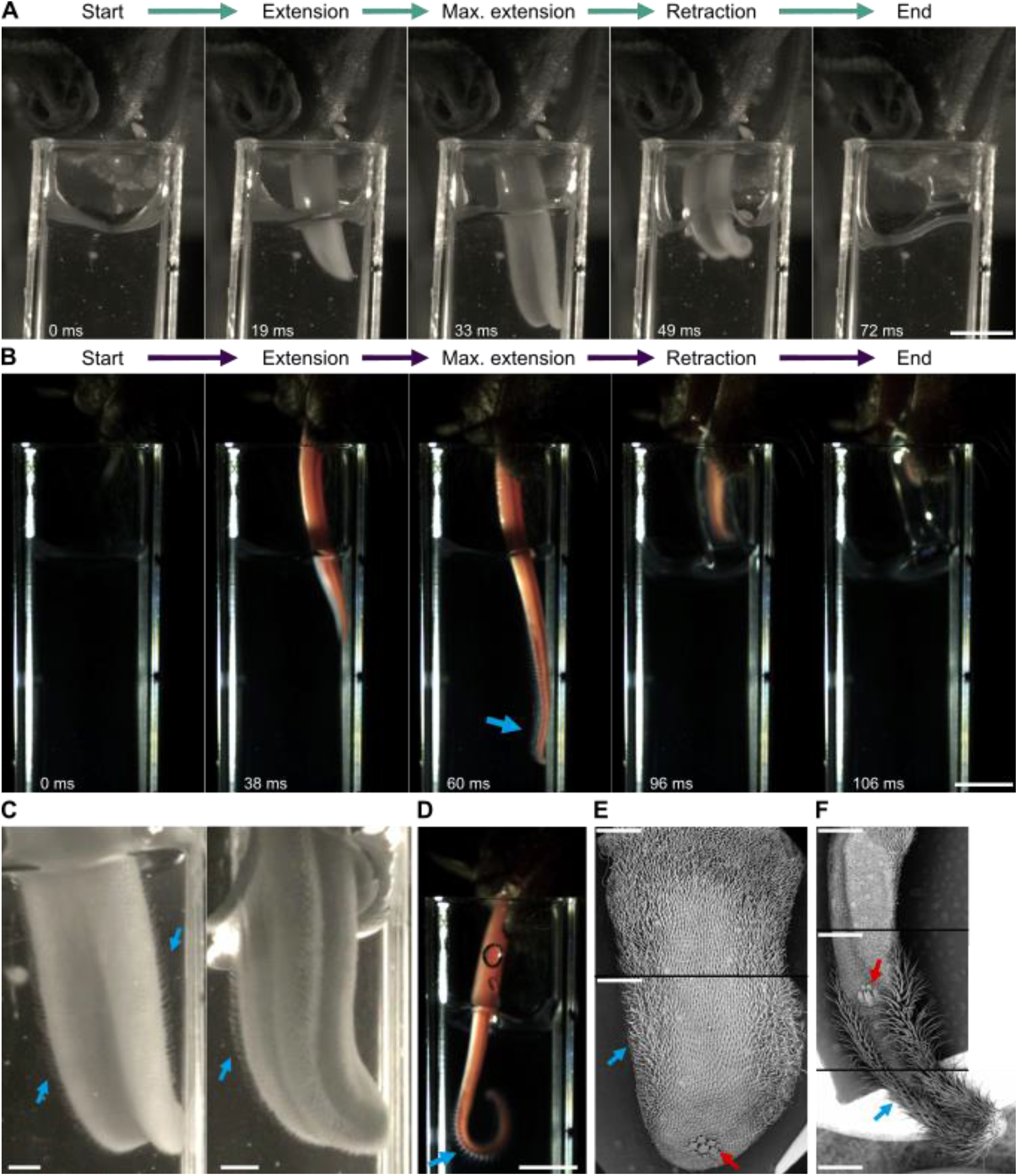
Lapping mechanisms and hair-like papillae of *Phyllostomus discolor* and *Anoura geoffroyi*. A) Frames from a high-speed free-flying video showing the nectar extraction sequence of a single lick in *P. discolor*. Time elapsed since the beginning of the cycle in milliseconds. Bar: 5 mm. B) Frames from a high-speed free-flying video showing the nectar extraction sequence of a single lick in *A. geoffroyi*. Time elapsed since the beginning of the cycle in milliseconds. Bar: 5 mm. Blue arrow points at hair-like papillae. C) Frames from a high-speed video showing the hair-like papillae (blue arrows) on the tongue of *P. discolor* in dorso-and ventro-lateral views. Bars: 1 mm. D). Frame from a high-speed video showing the hair-like papillae (blue arrow) on the tongue of *A. geoffroyi*. Bar: 5 mm. E) Scanning electron micrograph of the tongue of *P. discolor*, dorsal view. Blue arrow points at hair-like papillae, red arrow points at horny papillae. Bars: 1 mm. F) Scanning electron micrograph of the tongue of *A. geoffroyi*, dorsal view. Blue arrow points at hair-like papillae, red arrow points at horny papillae. Bars: 1 mm.

An interesting difference between the dynamic tongue shapes between these species is that in *P. discolor* there is a channel formed in the center of the tongue on the dorsal surface. This trough is formed by muscular action that generates a local bending in the sagittal plane of the tongue (Fig. 1A). The tongue does not normally present a trough on the dorsal surface (Fig. 1E); this channel is formed as the tongue is pulled out during feeding. During most of the tongue protrusion, the medial trough is not visible, it progressively becomes longer and deeper near maximum protrusion, and persists during retraction (Fig. 1A). Additionally, in the lateral part of the tongue of *P. discolor*, we could see dilation of blood vessels as the tongue was extended (Fig. S1). In some cases, *P. discolor* individuals took nectar from the nectar surface (instead of fully submerging the tongue), bending and folding the tongue before barely touching the surface, this was more common in hand-held videos and sometimes at the beginning of some free-flying visits.

We observed hair-like papillae extending from the lateral surface of the tongue of *P. discolor* individuals during nectar extraction (Fig. 1C). Even longer hair-like papillae were also observed in the high-speed videos of *A. geoffroyi* during feeding (Fig. 1D). The tongues of both species show hydrophilic properties; when the tongue meets the nectar and retracts, a layer of liquid adheres to the tongue on the dorsal and lateral sides, which is then transported into the mouth (Fig. S2). The hair-like papillae of *P. discolor* are located along the sides of the tongue more proximally than the horny papillae observed near the tip of the tongue (Fig. 1E), and they become longer and more abundant towards the middle section of the tongue (Fig 1E). This longer hair- like papillae will be potentially more exposed during formation of the ventro-caudal hook by the bending of the tip.

In *A. geoffroyi*, hair-like papillae are also present along the tongue sides but more distally than the horny papillae, covering the lingual surface all the way to the tongue tip, where they are highly abundant (Fig 1F). The hair-like papillae of *P. discolor* are shorter and less abundant than those of *A. geoffroyi* (see Quinche et al., 2023 for a deeper discussion of the lingual papillae of *P. discolor*). We observed 6 types of papillae in *A.geoffroyi*, 4 mechanical papillae: hair-like filiform papillae (Hlf), horny papillae (Hn), single-pointed filiform papillae (Spf), and branched filiform papillae (Brf); and 2 gustatory papillae: circumvallate papillae (Cv) and fungiform papillae (Fig. 2).

**Fig. 2.**
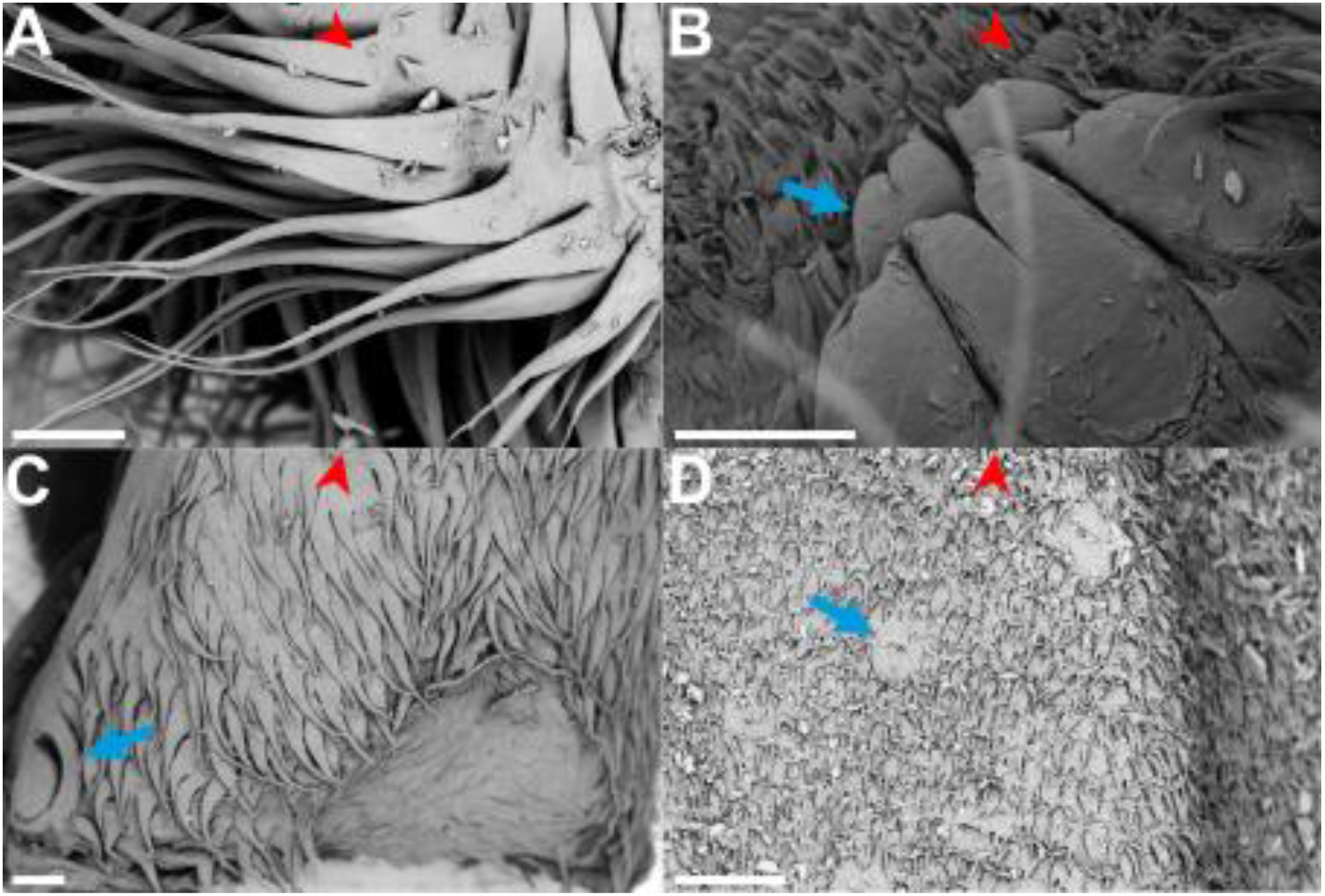
Scanning electron micrograph of the dorsal tongue papillae of *Anoura geoffroyi*. The red arrowheads point to the tip of the tongue. (A) Hair-like papillae (Hlf). (B) Horny papillae (Hn) (blue arrow) surrounded by branched filiform papillae (Brf). (C) Single-pointed filiform papillae (Spf) and circumvallate papillae (Cv) (blue arrow). (D) Fungiform papillae (Fg) (blue arrow) surrounded by branched filiform papillae. Bar 200 µm.

Although both species exhibit similar (potentially convergent) hair-like papillae and tongue kinematics, there are differences between the two lapping mechanisms, as exemplified in the plot of tongue reciprocation patterns (Fig. 3). *P. discolor* used more licks per visit, and the number of licks was more variable (4–77 licks) than in *A. geoffroyi* (1–7 licks) (Fig. 3A). Also, *P. discolor* continued feeding in most visits until it was unable to reach more nectar, making longer visits than *A. geoffroyi* (Figs. 3A, 4B). Furthermore, the insertion of the snout into the test container was constant throughout the first lick cycle in *P. discolor*, in contrast to *A. geoffroyi*, which inserted the snout more towards the end of the first lick (Fig. 3B). Additionally, we observed no unsuccessful attempts by *A. geoffroyi* to access nectar, unlike *P. discolor*. As expected, tongue tip insertion in *A. geoffroyi* was greater than that in *P. discolor* (Fig. 3B). The duration of protrusion and retraction in both species was similar, with both species reaching maximum tongue protrusion at around 50% of the lick (Fig. 3B).

**Fig. 3.**
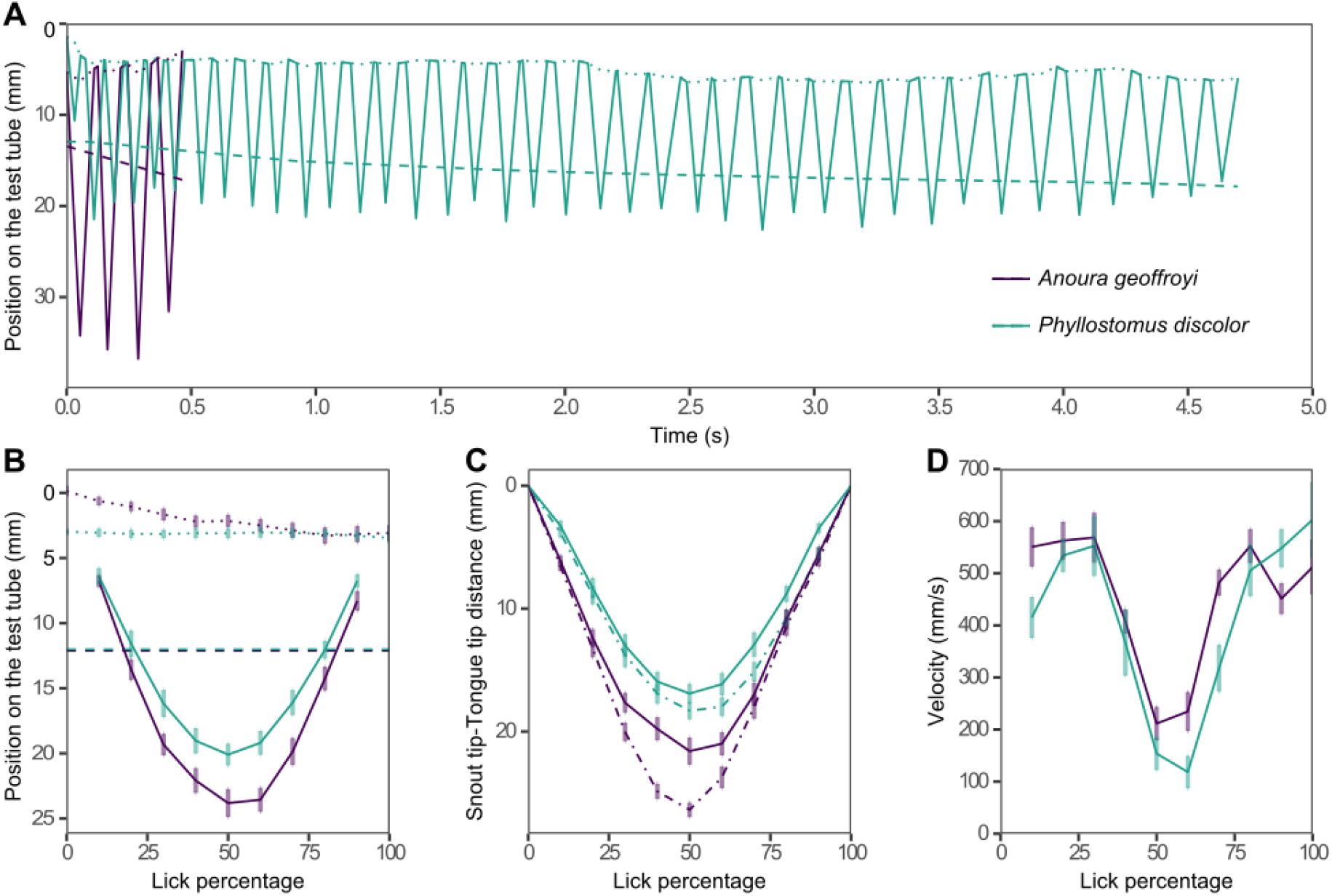
Tongue movement pattern and kinematics for *P. discolor* (green) and *A. geoffroyi* (purple). (A) Position on the test container of snout tip (dotted line), tongue tip (solid line), and nectar depth (dashed line) for one complete visit through time. (B) Position on the test container of snout tip (dotted line), tongue tip (solid line), and nectar depth (dashed line) for the first lick of multiple visits as a function of time (shown as percent of the lick). (C) Distance between the snout tip and the tongue tip in the y-axis (solid line) and following the curvature of the tongue (dot-dash line) for the first lick of multiple visits as a function of time (shown as percent of the lick). (D) Velocity of the tongue tip for the first lick of multiple visits as a function of time (shown as percent of the lick). In (A), (B) and (C), we show the Means ± s.e of the first lick of 9 visits for *P. discolor* and 17 for *A. geoffroyi*.

**Fig. 4.**
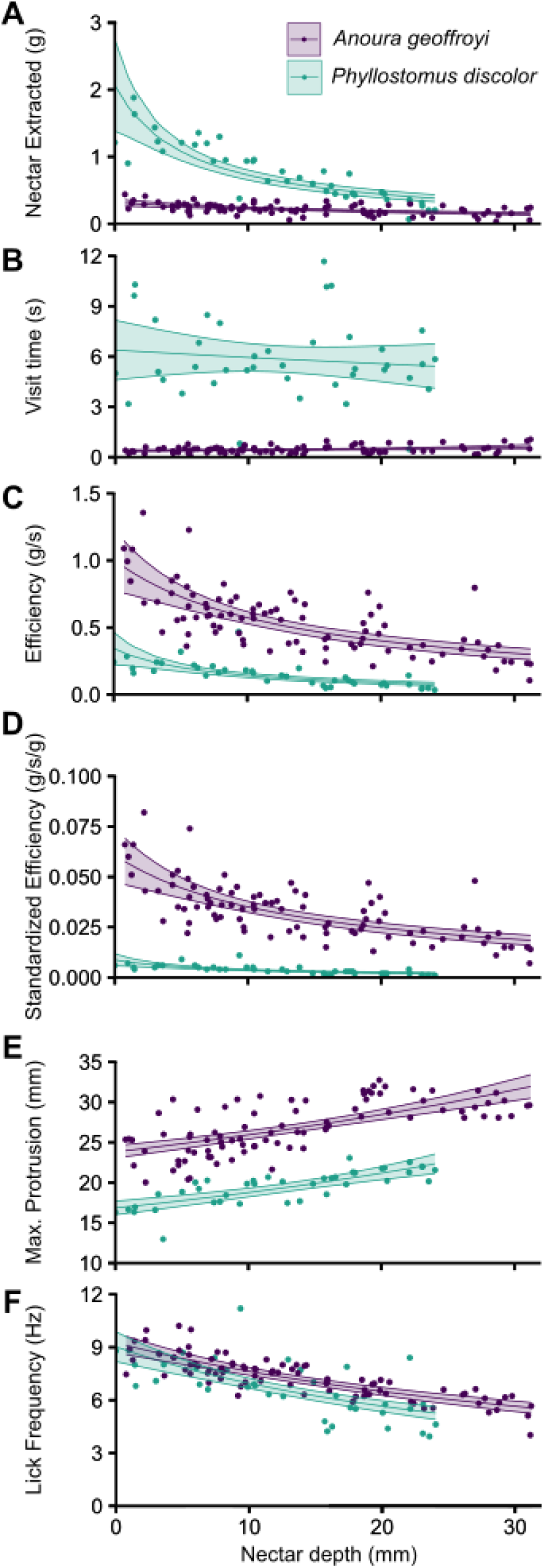
Drinking behavior of *P. discolor* and *A. geoffroyi*. Depth of nectar depth (mm) vs. (A) Nectar extracted per visit (g), (B) Visit time (s), (C) Feeding efficiency (g/s) (D) Size-adjusted (standardized) efficiency (g/s/g), (E) Maximum tongue protrusion (mm), and (F) Lick frequency (Hz) of the complete visits. Individual points represent different visits. The trend line represents the GLM between the couple of variables. The shaded gray area represents the 95% confidence interval of the model. 40 visits for *P. discolor* and 88 visits for *A.geoffroyi*.

Figure 3C represents the curvature of the tongue as the difference between the total protrusion of the tongue (dotted line) and the straight distance between the tongue tip and snout tip (solid line). Both species curved their tongues during feeding and reached maximum curvature around the maximum tongue protrusion (around 50% of the cycle). In general, *A. geoffroyi* curved its tongue more than *P. discolor*. In addition, the pattern of curvature throughout the cycle was different for both species: *P. discolor* curved the tongue from 30% to the end of the lick, retracting the curved tongue towards the mouth, whereas *A. geoffroyi* curved it from approximately 20% to 60% of the lick, retracting the tongue straight towards the mouth (Fig. 3C). Finally, figure 3D shows that the pattern of change in velocity during protrusion was similar for both species; however, *A. geoffroyi* showed a higher retraction velocity than *P. discolor* (Fig. 3D). The change in tongue velocity during protrusion and retraction (slope) was similar for both species, indicating a symmetrical pattern of tongue movement during feeding (Fig. 3D).

### Drinking behavior

To compare different metrics of nectar feeding behavior, we extracted data from 40 visits of five *P. discolor* individuals and 88 visits of 11 *A. geoffroyi* individuals at different nectar depths. Both explanatory variables, species and nectar depth, had a significant effect on the amount of nectar extracted, but not their interaction (Table 1). The amount of nectar extracted per visit decreased with increasing nectar depth for both *A. geoffroyi* and *P. discolor* (Fig. 4A). Nectar extracted by *P. discolor* was initially higher but decreased faster with increasing nectar depth than that extracted by *A. geoffroyi*. We found significant effects of species and nectar depth on the visit time, as well as their interaction (Table 1). At all nectar depths, *A. geoffroyi* visited the feeder much more quickly than *P. discolor* (Fig. 4B). For *A. geoffroyi*, there is a trend of increasing visit time with depth; however, while there are significant differences among depths for P. discolor, there is no clear trend of increasing or decreasing visit time. Species, nectar depth, and their interaction also significantly affected size-adjusted efficiency (Table 1). The efficiency and size- adjusted (standardized) efficiency decreased significantly towards deeper nectar depth for both species (Fig. 4C,D). However, this decrease was more pronounced for *P. discolor* than for *A. geoffroyi*. Nectar depth, species, and their interaction significantly affected maximum tongue protrusion (Fig. 4E, Table 1). The mean maximum tongue protrusion was higher in *A. geoffroyi* than in *P. discolor* at all nectar depths (Fig. 4E). It’s important to note that we did not experimentally push *A. geoffroyi* to its maximum tongue extension during feeding (i.e., we did not continue testing its ability to reach deeper nectar until it could no longer do so). Therefore, its maximum tongue length could potentially exceed the length reported here. Nectar depth had a significant effect on lick frequency, however, species did not (Fig. 4F, Table 1). The lick frequency of the complete visits was similar between both species, although lower (but again without reaching significance) at almost all depths for *P. discolor*.

**Table 1.**
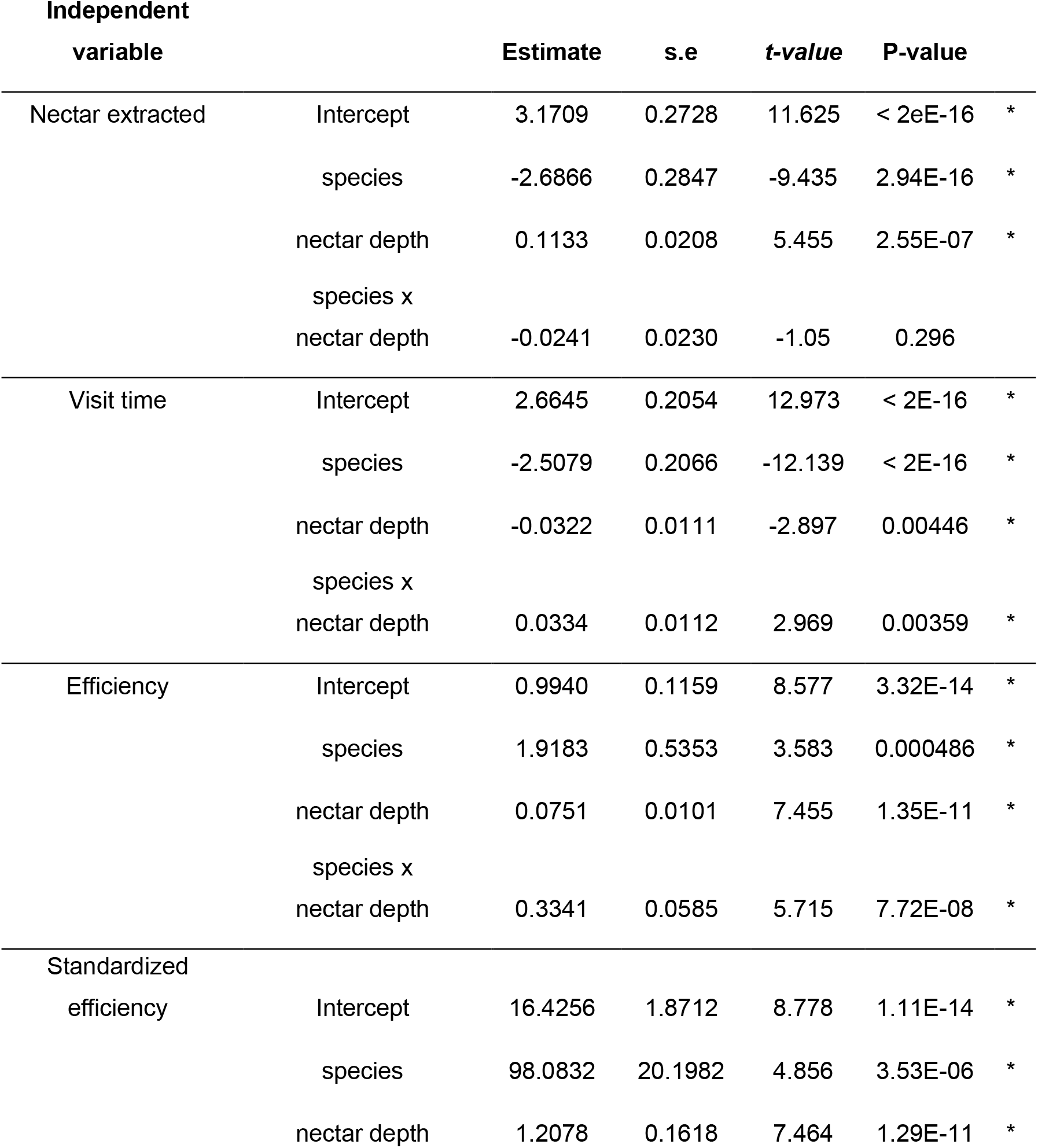
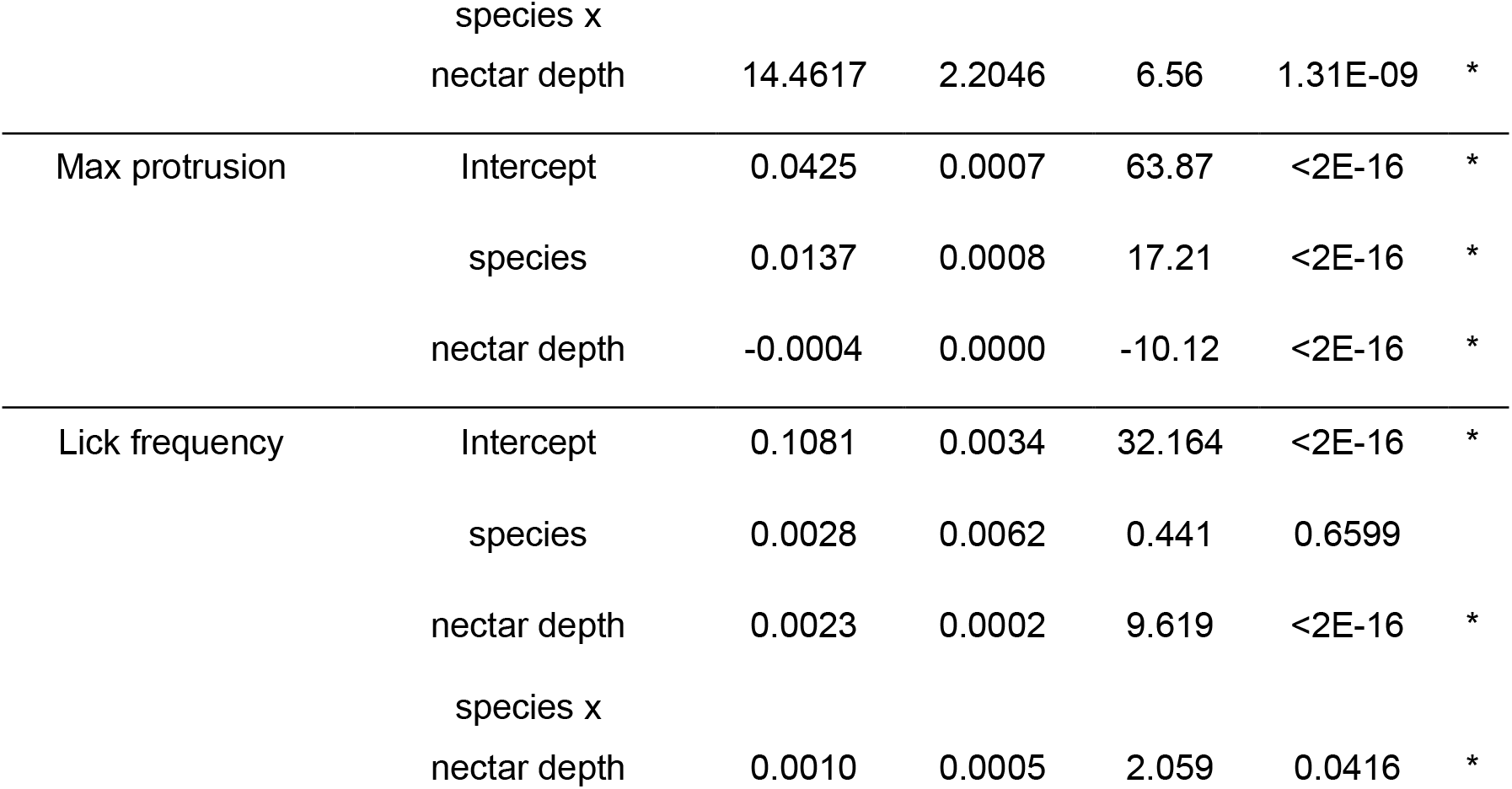
Generalized Linear Model (GLM) results for drinking behavior metrics and the effect of species, nectar depth, and their interaction.

| Independent variable |  | Estimate | s.e | t-value | P-value |  |
| --- | --- | --- | --- | --- | --- | --- |
| Nectar extracted | Intercept | 3.1709 | 0.2728 | 11.625 | < 2eE-16 | * |
|  | species | -2.6866 | 0.2847 | -9.435 | 2.94E-16 | * |
|  | nectar depth | 0.1133 | 0.0208 | 5.455 | 2.55E-07 | * |
|  | species x nectar depth | -0.0241 | 0.0230 | -1.05 | 0.296 |  |
| Visit time | Intercept | 2.6645 | 0.2054 | 12.973 | < 2E-16 | * |
|  | species | -2.5079 | 0.2066 | -12.139 | < 2E-16 | * |
|  | nectar depth | -0.0322 | 0.0111 | -2.897 | 0.00446 | * |
|  | species x nectar depth | 0.0334 | 0.0112 | 2.969 | 0.00359 | * |
| Efficiency | Intercept | 0.9940 | 0.1159 | 8.577 | 3.32E-14 | * |
|  | species | 1.9183 | 0.5353 | 3.583 | 0.000486 | * |
|  | nectar depth | 0.0751 | 0.0101 | 7.455 | 1.35E-11 | * |
|  | species x nectar depth | 0.3341 | 0.0585 | 5.715 | 7.72E-08 | * |
| Standardized efficiency | Intercept | 16.4256 | 1.8712 | 8.778 | 1.11E-14 | * |
|  | species | 98.0832 | 20.1982 | 4.856 | 3.53E-06 | * |
|  | nectar depth | 1.2078 | 0.1618 | 7.464 | 1.29E-11 | * |
|  | species x<br>nectar depth | 14.4617 | 2.2046 | 6.56 | 1.31E-09 | * |
| Max protrusion | Intercept | 0.0425 | 0.0007 | 63.87 | <2E-16 | * |
|  | species | 0.0137 | 0.0008 | 17.21 | <2E-16 | * |
|  | nectar depth | -0.0004 | 0.0000 | -10.12 | <2E-16 | * |
| Lick frequency | Intercept | 0.1081 | 0.0034 | 32.164 | <2E-16 | * |
|  | species | 0.0028 | 0.0062 | 0.441 | 0.6599 |  |
|  | nectar depth | 0.0023 | 0.0002 | 9.619 | <2E-16 | * |
|  | species x<br>nectar depth | 0.0010 | 0.0005 | 2.059 | 0.0416 | * |

## DISCUSSION

### Feeding mechanism and associated morphology

For the first time, we investigated the nectar extraction mechanism in two bat species with differing levels of specialization for a nectarivorous diet, *Phyllostomus discolor* and *Anoura geoffroyi*. Both species employed a lapping technique, characterized by stereotypic reciprocating movements of their long and highly mobile tongues (Figs. 1A,B, Movies 1, 2 in supplementary material). The tongues of *P. discolor* and *A. geoffroyi* possess hair-like papillae that increase surface area for liquid coating via adhesion, and thus function in this brush-tongue lapping technique, a widespread mechanism among nectar-feeding animals (Cuban et al., 2022; Harper et al., 2013; Hewes et al., 2023; Richardson and Wooller, 1986; Yang et al., 2014). However, differences in the relative positions and structures of these papillae (Fig. 1E,F) suggest distinct anatomical origins and analogous adaptations for nectar collection. Despite these differences, the shared lapping mechanism highlights convergence in their general nectar-feeding strategies.

In Glossophaginae bats, nectar is trapped between the hair-like papillae on the tongue during each lap (Harper et al., 2013). For *P. discolor*, the tongue’s backward bending and central folding create a channel that increases nectar capture by optimizing the brush-like papillae’s contact with nectar. This curving mechanism, observed in both species, likely enhances their ability to ’mop’ nectar inside flowers, maximizing extraction efficiency. Although both species share the lapping mechanism, *P. discolor* exhibited less tongue insertion and curvature but more snout insertion and licks per visit compared to *A. geoffroyi* (Fig. 3B,C). These differences reflect *P. discolor*’s lesser morphological and behavioral specialization for nectar feeding. Its shorter and thicker tongue (Figs. 1E,F, 3B), limits its ability to reach greater nectar depths or achieve more pronounced tongue curvature (Nicolay and Winter, 2006). This contrasts with the specialized tongue morphology of *A. geoffroyi*, which aligns with adaptations seen across Glossophaginae species (Harper et al., 2013; Muchhala et al., 2024; Nicolay and Winter, 2006).

Foraging behavior influenced some of the feeding mechanism characteristics. Landing in *P. discolor* and hovering in glossophagines (e.g., *A. geoffroyi*), relate to the resulting differences in snout insertion and the number of licks per visit (Fig. 3B). Hovering imposes energetic constraints on glossophagines due to its high energy demand (Winter, 1998), limiting visit duration. In contrast, *P. discolor* faces no such constraint, enabling longer visits with more licks. Additionally, *P. discolor* initiated visits after inserting its snout, whereas *A. geoffroyi* progressively inserted its snout, enhancing its ability to locate nectar efficiently. Together with the fact that, unlike *P. discolor*, we did not observe any unsuccessful attempts by *A. geoffroyi* to access nectar, we suggest that *A. geoffroyi* is better at locating nectar than *P. discolor*. Some studies have highlighted the ability of specialized nectar bats to effectively detect and locate nectar sources due to sensory adaptations (Amichai et al., 2023; Gonzalez-Terrazas et al., 2016).

We observed similarities between the two species in tongue protrusion and retraction duration, as well as tongue velocity during one licking cycle (Fig. 3D). Tongue reciprocation followed a symmetrical path, with both species exhibiting comparable velocity changes during protrusion and retraction. However, *A. geoffroyi* retracted the tongue faster than *P. discolor*, likely due to specialized vascular and muscular adaptations to feed on nectar in Glossophaginae species (Griffiths, 1978; Harper et al., 2013). Further studies are needed to understand the muscular adaptations in *P. discolor*, which achieved lick frequencies comparable to the specialized *A. geoffroyi*.

### Drinking behavior

All evaluated feeding behavior variables differed significantly between species, except for licking rate. Nectar depth influenced all drinking behavior variables (Table 1). Both species extracted less nectar as depth increased, consistent with previous findings on nectar-feeding bats (Gonzalez-Terrazas et al., 2012; Nicolay and Winter, 2006). This limitation arises from the lapping mechanism, where nectar adheres to the tongue’s surface and is trapped between papillae (Harper et al., 2013). The extent of tongue submersion directly affects nectar extraction (Crompton and Musinsky, 2011; Reis et al., 2010).

Interestingly, *P. discolor* extracted more nectar per visit than *A. geoffroyi* (Fig. 4A), despite the latter’s specialized tongue morphology. This difference is attributed to *P. discolor*’s foraging style (i.e. landing), which allows it to forage on flowers for a longer period of time, which often involves draining flowers (Fischer, 1992; Heithaus et al., 1974; Pedrozo et al., 2018). This strategy may be advantageous but comes with increased predation risks (Nicolay and Winter, 2006) and requires robust plant structures to support its weight. In addition, their shorter and less abundant hair-like papillae (Quinche et al., 2023) would potentially trap less nectar compared to the more abundant and longer papillae in *A. geoffroyi* (Figs. 1E,F, 2A) (Nasto et al., 2018), especially when extracting the last bits of nectar deep inside flowers (where the longer distal hair-like papillae of *A. geoffroyi* would be able to reach better).

The differences in foraging behavior between the two bat species significantly influenced their visitation times (Figs. 1A, 4B). In our experiments, *P. discolor* individuals remained perched and often continued feeding until it became challenging to access the remaining nectar. In contrast, the shorter visitation times of *A. geoffroyi* are likely tied to the energetic demands of hovering during feeding (Winter, 1998). While the presence of a landing structure in our experimental setup may have extended the visit duration of *P. discolor*, making it longer than previously reported (Nicolay and Winter, 2006), we posit that our findings align with the species’ natural foraging behavior on flowers that provide such structures (Gribel et al., 1999; Hopkins, 1984; Pedrozo et al., 2018; Sazima and Sazima, 1977).

Nectar extraction efficiency, both standardized and non-standardized by weight, was significantly lower in P. discolor compared to *A. geoffroyi* (Fig. 4C,D), consistent with previous findings comparing glossophagine and non-glossophagine species (Nicolay and Winter, 2006). This difference in efficiency is influenced by the varying degrees of morphological specialization for nectar feeding between the species, including tongue length and the size, and density of hair-like papillae (Fig. 1E,F). Additionally, the feeding mode—landing in *P. discolor* versus hovering in *A. geoffroyi*—plays a significant role, highlighting the combined impact of morphological and behavioral traits on nectar extraction efficiency (Nicolay and Winter, 2006). Similar to earlier studies, our results showed that nectar depth has a significant negative effect on the extraction efficiency of *P. discolor* (Table 1). However, the efficiency observed for *P. discolor* in our experiments was lower than reported in previous studies (Nicolay and Winter, 2006), likely due to the availability of a landing structure in our setup. While the landing strategy in our experiments may result in reduced feeding efficiency, it likely lowers the energetic costs of feeding compared to hovering (Voigt, 2004). Nonetheless, energetic costs associated with landing and take-off also need to be accounted for when comparing the overall energetics of these feeding strategies.

Although dietary specialization enables species to exploit specific resources more efficiently, it comes with a tradeoff: specialization can limit the ability to utilize alternative resources when preferred food is scarce (Rex et al., 2010). For *P. discolor*, the ability to switch to alternative food resources has been documented in various diet studies (Bonaccorso, 1979; Kwiecinski, 2006). The morphological, biomechanical, and behavioral differences between *P. discolor* and *A. geoffroyi* significantly influence the types of resources they can access and feed on effectively (Freeman, 1995; Gonzalez-Terrazas et al., 2012; Winter and von Helversen, 2003). The foraging strategy and tongue morphology of each species largely determine and restrict the types of flowers and other food items they can exploit. For example, the landing strategy and larger body size of *P. discolor* limit its access to smaller or more delicate flowers that cannot support its weight (Fischer, 1992; Gonzalez-Terrazas et al., 2012). Moreover, *P. discolor* is better suited for flowers with large openings and shorter corollas, as it relies more on snout insertion and possesses a wider, shorter, and less flexible tongue compared to *A. geoffroyi* (Figs. 1E,F, 2B,C). These phenotypic differences, coupled with the omnivorous diet of *P. discolor*, allow the two species to exploit different resources and coexist, even when their primary food source—nectar—overlaps (Finke and Snyder, 2008).

### Conclusions and future directions

Our study highlights convergence in the nectar extraction mechanisms and associated morphology of the omnivorous *P. discolor* and specialized nectar- feeder *A. geoffroyi*. Both species employ an equally speedy brush-tongue lapping technique facilitated by the exposure of hair-like papillae. Notably, *P. discolor* demonstrates tongue mobility comparable to that of *A. geoffroyi*, as evidenced by their similar lick frequencies. Furthermore, *P. discolor* exhibits a combination of morphological and behavioral adaptations that enable efficient nectar exploitation while retaining the flexibility to feed on alternative resources. This underscores the versatility of *P. discolor* and its capacity to adapt to diverse dietary niches.

Although *P. discolor* employs the same general feeding mechanism as *A. geoffroyi* and other Glossophaginae species, it remains unclear whether this mechanism is driven by the hemodynamic erection of papillae, as observed in a Glossophaginae species (Harper et al., 2013). Future research should investigate this possibility, as well as determine the energetic costs associated with its feeding behavior and compare these with other *Phyllostomus* species and nectarivorous bats. Exploring the trade-offs within and between traits involved in food acquisition could further illuminate the evolutionary shifts in feeding modes and levels of specialization among nectar-feeding bats.

## LIST OF SYMBOLS AND ABBREVIATIONS

Hlf: hair-like filiform papillae
Hn: horny papillae
Brf: branched filiform papillae
Spf: single-pointed filiform papillae
Fg: fungiform papillae
Cv: circumvallate papillae

## ACKNOWLEDGEMENTS

We thank the members of the Behavioral Ecophysics and Santana labs at the University of Washington for their invaluable discussions and support. We are grateful to everyone at the Colibrí Gorriazul Research Center (Centro de Investigación Colibrí Gorriazul - CICG) in Colombia, Francisco Javier Urrea, and Camila Valdés Cardona for their assistance during the field seasons. We thank Hacienda La Cabaña for allowing us to conduct fieldwork at their facilities. Special thanks to Lucero Rojas, Parmenio Simbaqueba, and Mary Simbaqueba for making CICG an enjoyable place to conduct research. Additional thanks to Rafael Quinche and Nathaly Garzón for their help with fieldwork.

## FOOTNOTES

### Author contributions

Conceptualization: L.L.Q., A.R.-G; Methodology: L.L.Q., A.R.-G., F.G.-A.; Formal Analysis: L.L.Q.; Investigation: L.L.Q., F.G.-A.; Resources: A.R.-G.; Writing original draft: L.L.Q., A.R.-G., H.F.L.-A.; Writing –review & editing: L.L.Q., A.R.-G., F.G.-A., S.E.S.; Visualization: L.L.Q., F.G.- A.; Supervision: L.L.Q., A.R.-G.; Project administration: A.R.-G., S.E.S., H.F.L.-A.; Funding acquisition: A.R.-G., S.E.S., H.F.L.-A.

### Data availability

All the necessary data are reported within the manuscript and supplementary information. Further supportive data or clarifications are available from the corresponding author upon reasonable request.

## SUPPLEMENTARY INFORMATION

**Movie 1. Highspeed video of *Phyllostomus discolor* feeding on nectar.** At maximum tongue extension during each lick, the tongue folds medially, and hair-like papillae are erected. Recording rate=1000 fps; playback speed=20 fps.

**Movie 2. Highspeed video of *Anoura geoffroyi* feeding on nectar.** Note how hair-like papillae are erected and the tongue curves in multiple directions. Recording rate=500 fps; playback speed=12.5 fps.

**Fig. S1.**
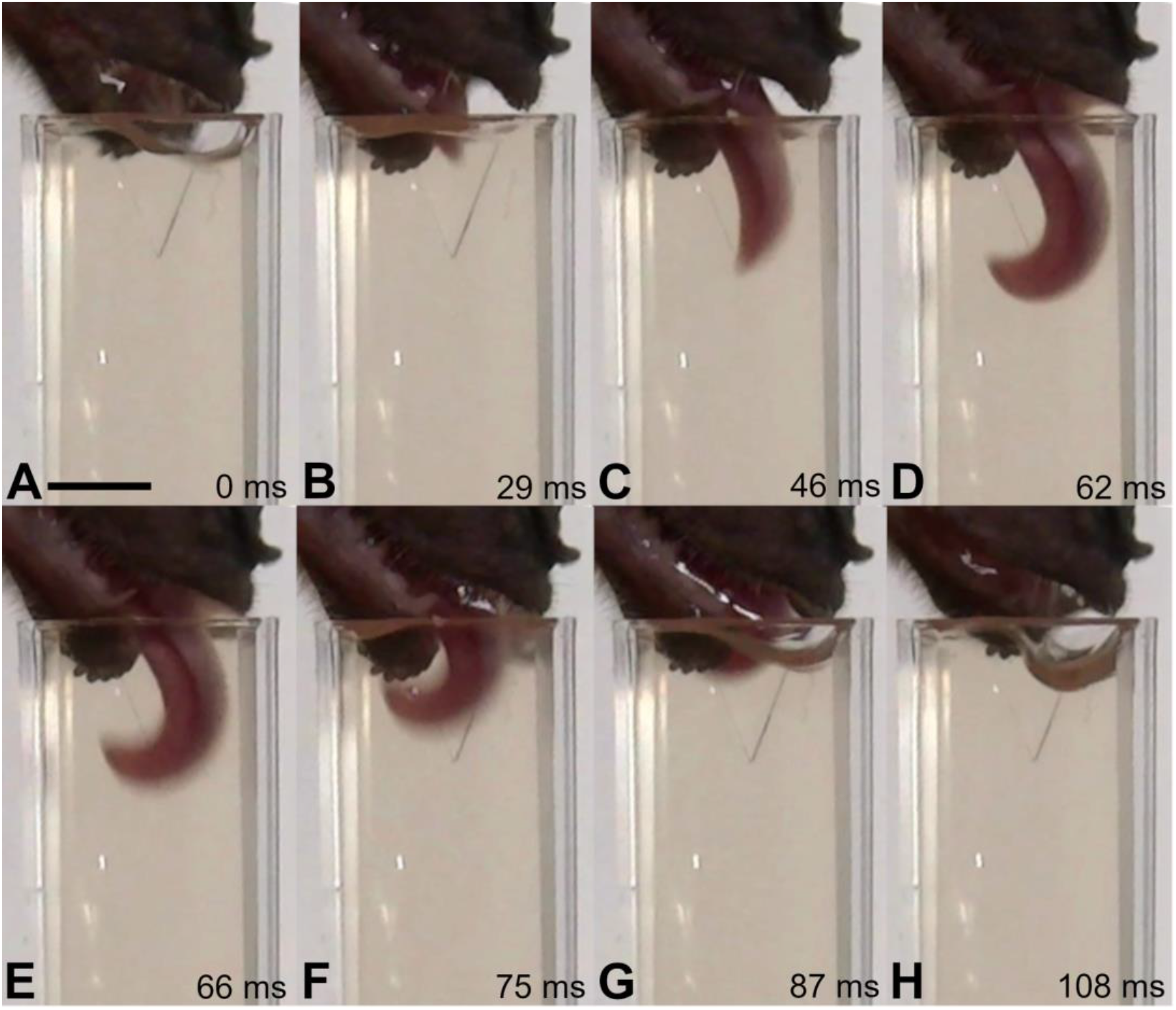
Nectar extraction sequence of a single lick in *Phyllostomus discolor* (lateral view), illustrating the tongue’s curvature into a ’hook’ shape. Frames from a high-speed hand-held video. (A) start of the cycle. Bar: 5 mm. (B) and (C) protrusion. (D) maximum protrusion. (E), (F), and (G) retraction. (H) end of the cycle. Time elapsed since the beginning of the cycle in milliseconds.

**Fig. S2.**
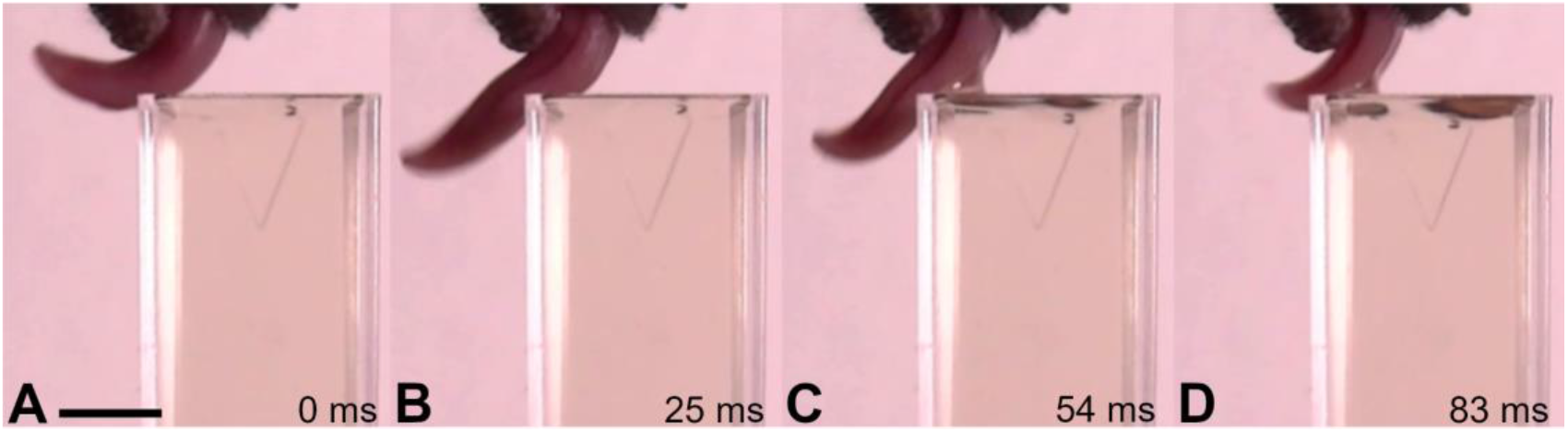
Sequence of nectar extraction in *Phyllostomus discolor* (lateral view), illustrating the tongue’s hydrophilic properties. Frames from a high-speed hand-held video. (A) protrusion. Bar: 5mm. (B) maximum protrusion. The lateral line of blood vessel dilatation is much more evident (C) and (D) retraction, nectar layer is forming on the dorsal part of the tongue. Time elapsed since the beginning of the cycle in milliseconds.

